# *In vivo* evolution of *Candida auris* multi-drug resistance in a patient receiving antifungal treatment

**DOI:** 10.1101/2025.05.13.653818

**Authors:** Tristan W. Wang, Nicole Putnam, J. Kristie Johnson, Mary Ann Jabra-Rizk

## Abstract

*Candida auris* is associated with life-threatening invasive disease due to high level of drug resistance. We present a clinical case of *C. auris* multi-drug resistance development in a single patient acquired during antifungal treatment. Five isolates were prospectively recovered from a transplant patient receiving antifungal therapy over a one-year period. While isolates were initially only resistant to fluconazole, the terminal isolate became resistant to caspofungin and amphotericin B with significant increase in micafungin MIC. Sequencing of *ERG11* and *FKS1* genes identified mutations associated with fluconazole and echinocandin resistance in the multi-drug-resistant isolate, underscoring the threat of therapy-induced development of resistance.

## BACKGROUND

*Candida auris* is a fungal species that has emerged as a nosocomial pathogen causing outbreaks of life-threatening invasive disease worldwide [1]. Most concerning about *C. auris* is the unprecedented level of drug resistance and the ability to develop resistance to all classes of antifungals, severely limiting treatment options [2]. This multidrug-resistant (MDR) pattern has been observed in around 40% of clinical isolates, and outcomes of infection have generally been poor with mortality rates approaching 68% [3]. Due to its high transmissibility and prevalent antifungal resistance, *C. auris* is categorized as a critical priority pathogen by the World Health Organization [4].

Studies of antifungal susceptibility profiles on *C. auris* isolates revealed that up to 90% of isolates were resistant to fluconazole, 8% to amphotericin B and 2% to echinocandins with the South Asia clade I exhibiting the greatest percentage of resistant isolates [5]. The echinocandins are currently the antifungal drugs of choice for treatment of *C. auris* infection due to limited side effects and the low rate of resistance. However, studies reporting development of echinocandin resistance during antifungal therapy are on the rise, primarily associated with *FKS1* gene mutations [6]. Another unique feature reported in some clinical *C. auris* isolates is cell aggregation, with aggregative and non-aggregative phenotypes exhibiting different biofilm forming abilities, drug susceptibility profiles and transcriptional changes induced by antifungal exposure [7, 8].

In this study, we present a case where *C. auris* isolates from a patient receiving antifungal treatment acquired resistance to caspofungin and amphotericin B during therapy. Prospectively recovered isolates were evaluated for antifungal susceptibilities and sequencing was performed to identify clade and the mechanisms of resistance. Furthermore, the isolates were phenotypically evaluated for fitness, ability to aggregate, and form biofilms.

## METHODS

### Patient case, clinical isolates and antifungal susceptibility

The study was approved by the University of Maryland, Baltimore Institutional Review Board (IRB, HP-00110276) and Institutional Biosafety Committee (IBC-00007835). Clinical samples were obtained as part of routine patient care and diagnostic workup. *C. auris* was identified in the clinical microbiology laboratory by matrix-assisted laser desorption/ionization time-of-flight mass spectrometry (MALDI-TOF MS) using the VITEK MS (bioMérieux) and confirmed by ITS sequencing. The patient was a 68-year-old man admitted for bilateral orthotopic lung transplant (BOLT) and remained hospitalized for over 1 year due to complications. A total of 5 *C. auris* isolates were recovered from various sites before the patient expired on day 386 (Fig. 1).

**Figure 1.**
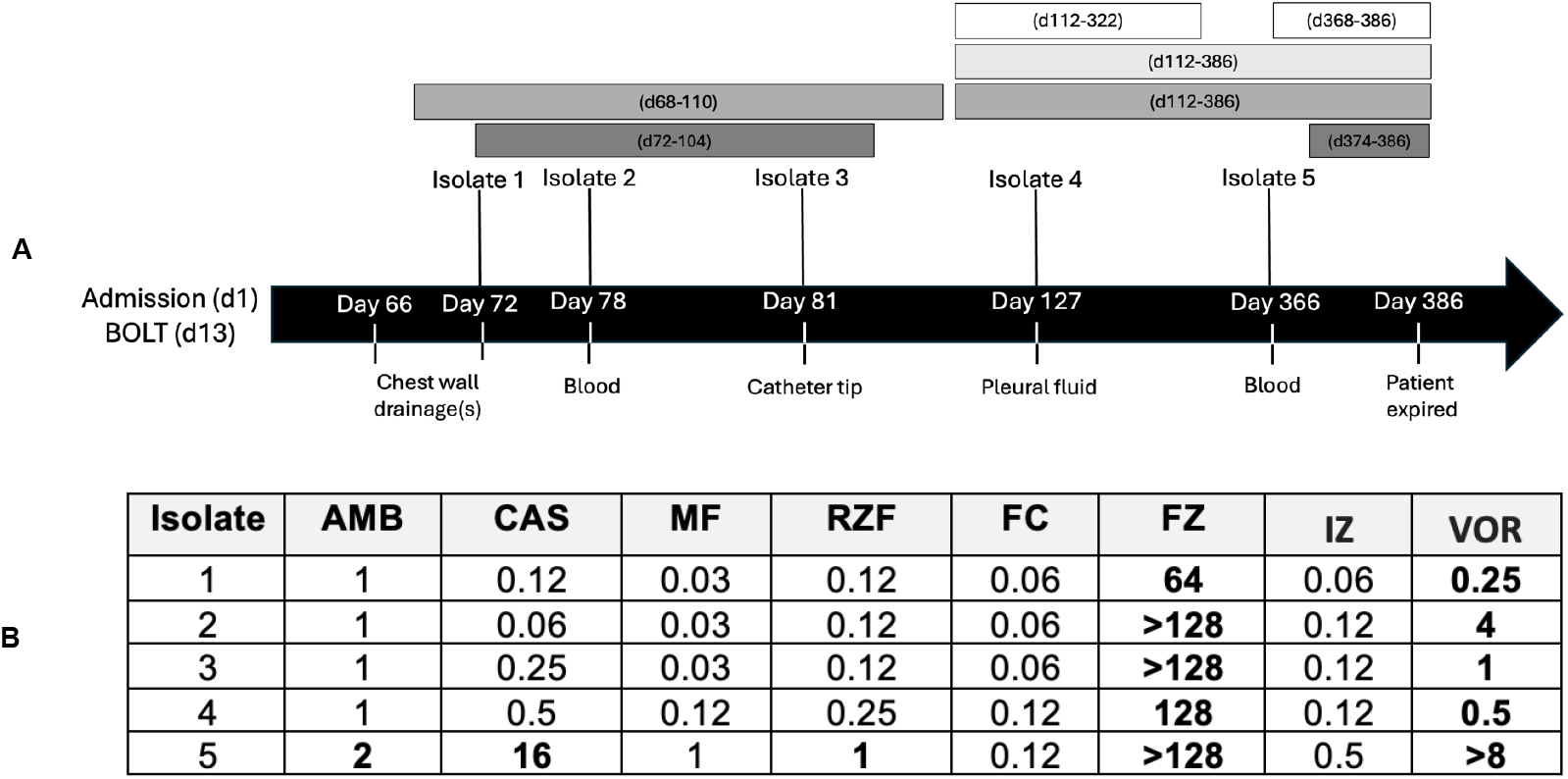
Clinical timeline of *Candida auris* isolation and antifungal treatment. **(A)** Patient was admitted for transplant evaluation with subsequent bilateral orthotopic lung transplant (BOLT). Timeline shows *C. auris* isolates from various locations and timepoints throughout hospitalization. Antifungals used for treatment: micafungin (medium gray); IV liposomal amphotericin B (dark gray). Antifungals for transplant prophylaxis: isavuconazole (white); inhaled amphotericin B (light gray). All isolates were resistant to fluconazole. **(B)** Clinical antifungal MIC (μg/ml) of recovered *C. auris* isolates. MICs shown in bold exceed tentative CDC MIC (μg/ml) breakpoints for resistance: fluconazole (FZ) ≥32; 5-flucytocine (FC) >8; amphotericin B (AMB) ≥2; caspofungin (CAS) ≥2; rezafungin (RZF) S ≤0.5; voriconazole (VOR) S ≤0.12; itraconazole (IZ) ≤0.125; micafungin (MF) ≥4; (https://www.cdc.gov/candida-auris/hcp/laboratories/antifungal-susceptibility-testing.html). There is no CDC advisory for FZ, IZ and VOR *breakpoints*; *MICs* are considered resistant based on breakpoints for other *Candida* species

### Antifungal susceptibility testing

Isolates were grown on yeast peptone dextrose agar (YPD) (Difco Laboratories) at 35°C. Isolated colonies were suspended in saline and used in antifungal susceptibility testing. To determine MICs the Sensititre YeastOne AST Plate (Thermo Scientific, YO4IVD) panels containing fluconazole, voriconazole, itraconazole, rezafungin, micafungin, caspofungin and 5-flucytosine were used. Amphotericin B was tested using the CLSI microbroth microdilution method [9]. MICs were read visually after 24h incubation at 35°C and resistance determined based on MICs exceeding tentative CDC breakpoints for *C. auris* [10].

### Growth kinetics

Isolates were grown overnight in YPD broth at 32°C. 200μl of 1×10^5^ cells/ml in YPD broth was inoculated in wells of 96-well flat-bottom microtiter plates and incubated at 37°C for 36h. Absorbance (OD_600_) was measured using a Cell Imaging Multi-Mode Reader (Cytation 5, Biotek).

### Comparative evaluation of biofilm formation and aggregation

Experiments were performed as we previously described [11]. Biofilm formation was assessed based on metabolic activity by seeding 200µl of 1×10^6^ cells/ml in RPMI1640 medium in wells of flat-bottom 96-well polystyrene microtiter plates. Following incubation at 37°C for 24h, wells were washed twice with PBS and biofilms evaluated using the MTS metabolic assay (Promega); color intensity was measured at OD_490_ using a microtiter plate reader. Aggregation was assessed based on formation of cell aggregates and sedimentation rate. Cell suspensions at 5×10^7^ cells/ml density were vigorously vortexed for 1min and formation of cell aggregates monitored and imaged by light microscopy. Sedimentation rate of aggregates was measured based on drop in absorbance readings at OD_600_ at 0 min and 30 mins post-vortexing using a Biomate 3S spectrophotometer (Thermo) and calculated as percent reduction in absorbance compared to initial reading.

### DNA Extraction, PCR, and Sequencing

Genomic DNA was extracted from clinical isolates and *C. auris* reference strain B8441 using the DNeasy Kit (Qiagen) following manufacturer’s instructions modified for yeast cultures. *ITS* and *RHA1* genes were sequenced to identify clade and the *ERG6, ERG11* and hotspots 1 and 2 in *FKS1* genes were sequenced to identify mutations. Amplification was performed using PCR Master Mix (Promega) and primers listed in Supp Table 1 following the method described by Carolus et al. [12]. Amplicons were purified using QIAquick PCR purification kit (Qiagen). Sanger sequencing was performed by GENEWIZ, Azenta Life Sciences. Mutations were identified by comparison to the *C. auris* reference strain B8441 (candidagenome.org).

### Stress sensitivity

Isolates (1×10^5^ cell/ml) were grown in 96-well microtiter plates (200μl/well) in the presence of SDS (0.12-3.5 mM), NaCl (0.15 mM-2.5 M) or H_2_O_2_ (2.5-40 mM) in YPD broth for 24h at 37°C and growth was measured based on absorbance readings at OD_600_ [13]. Cell suspensions (1×10^4^, 1×10^5^, 1×10^6^ cells/ml) were plated (10µl) on YPD agar supplemented with calcofluor white (10mg/L) and grown at 30°C for 24h and on YPD agar and incubated at 30°C, 37°C or 42°C or 24h. Isolates that showed growth at 42°C were considered thermotolerant.

### Data analysis

Statistical analysis of biofilm growth was performed using R statistical programming software. To compare differences among strains in sedimentation and *in vitro* biofilm forming capabilities, a one-way ANOVA with Tukey’s post host test was used. *P* values less than 0.05 were considered significant. Ggplot2 and ggpubr packages and Matplotlib library were used to construct figures.

## RESULTS

### Antifungal susceptibilities of *C. auris* clinical isolates

All 5 isolates were resistant to fluconazole and voriconazole. The terminal isolate (#5) became resistant to amphotericin B (MIC 2μg/ml), caspofungin (MIC >16 μg/ml), and rezafungin (MIC >1 μg/ml). Although within susceptible range, micafungin MIC for isolate 5 was significantly higher (0.03 to 1μg/m). All isolates were susceptible to 5-flucytosine and itraconazole (Fig. 1B).

### Sequencing and identification of clade and mutations

ITS sequencing identified all five isolates to belong to South Asia clade I. All isolates were found to have the Y132F mutation in the *ERG11* gene. Isolate four harbored the missense mutation M960I and the terminal MDR isolate resistant to caspofungin was found to harbor the F635Y mutation in hot-spot 1 of the *FKS1* gene. No mutations were identified in *FKS1* hot-spot 2 or the *ERG6* genes.

### Isolates exhibit difference in growth rates

Isolates had comparable growth rates up to 12h; subsequently, isolate #1 demonstrated highest growth rate and isolate #5 the lowest growth rate. Growth rates for isolates #1-3 were comparable at all timepoints (Fig. 2A).

**Figure 2.**
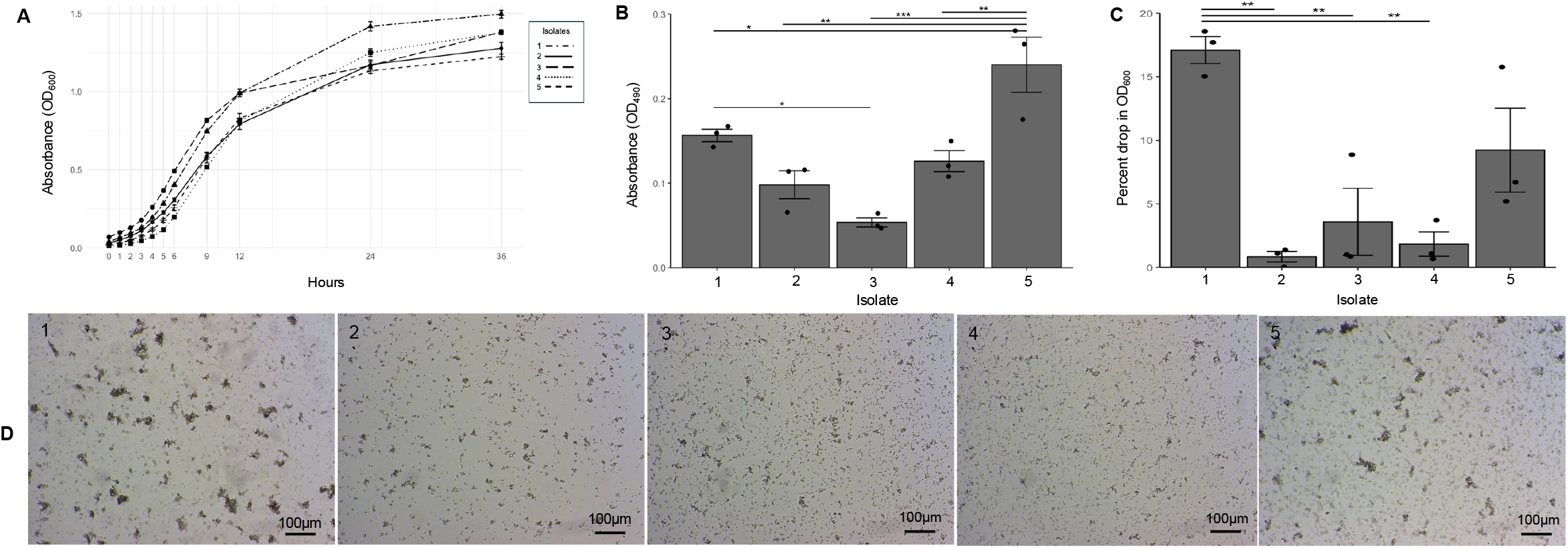
Comparative phenotypic evaluation of clinical isolates. **(A)** Evaluation of cell growth rates among isolates over 36h based on values of OD_600_. Line plots show mean and standard error of 10 technical replicates. **(B)** Measurement of metabolic activity of 24h biofilms based on values of OD_490_. Bar-plots show mean and standard error of mean of *n* = 3 biological replicates, each as an average of 4 technical replicates. **(C)** Measurement of rate of cell aggregate sedimentation over 30 mins. Bar-plots show mean (n = 3) of biological replicates, each an average of 3 technical replicates. **(D)** Representative light microscopy (10x) images of cell aggregates formed by isolates. Statistical analysis was performed by one-way ANOVA and post-hoc Tukey test with *P*-values representing significant differences. *0.01 < P ≤ 0.05, **0.001 < P ≤ 0.01, ***0.0001 < P ≤ 0.001. Non-significant differences are not shown.

### Isolates vary in biofilm forming abilities and aggregation

Based on metabolic activity, isolate #5 had the highest biofilm formation and #3 the lowest (Fig. 2B). Isolate #1 was the most aggregative followed by isolate #5 (Fig. 2C, D). No notable differences in aggregation for the other 3 isolates.

### Isolates vary in response to various stressors

No significant differences between the isolates were seen in their ability to grow in the presence of H_2_O_2_, SDS, NaCl and at 42°C. However, isolate #1 exhibited increased susceptibility to calcofluor white (Supp. Fig. 1).

## CONCLUSION

Greater than 90% of all *C. auris* clinical isolates are found to be resistant to fluconazole. Therefore, the high level of fluconazole resistance in our isolates was not surprising particularly as South Asia clade I isolates are overwhelmingly resistant to fluconazole [14]. The most commonly reported mechanism of resistance to the triazole antifungals is the acquisition of mutations in *ERG11*, the gene encoding lanosterol 14α-demethylase [15]. Nearly all fluconazole-resistant isolates of *C. auris* are found to have one of three mutations including Y132F which was present in all our 5 fluconazole resistant isolates [15].

Development of echinocandin resistance during treatment due to mutations in *FKS1* gene encoding 1,3-β-glucan synthase have been reported [6, 16]. In the terminal isolate resistant to echinocandins, we found the F635Y mutation in host-spot 1 previously shown to correlate with echinocandin resistance [17]. This mutation is of clinical significance as a study evaluating the impact of various mutations identified in *FKS1* in a disseminated infection mouse model, F635Y demonstrated the poorest response to caspofungin in terms of survival and organ fungal burden [17]. The only other mutation we found in the *FKS1* was a missense mutation (M690I) which was also previously identified in an *in vitro* evolved caspofungin resistant strain. However, the mutation did not seem to have a direct impact on susceptibility to caspofungin [18].

South Asia clade I isolates are variably resistant to amphotericin B and similar to echinocandins, resistance to amphotericin B was reported to develop in patients on therapy [14, 19, 20]. However, in line with findings from other studies, we did not identify nonsynonymous variants among amphotericin-susceptible and resistant isolates. A mutation in *ERG6* was found to be associated with increased resistance to amphotericin B [20], but was not found in the current study. While unknown genetic elements may be driving amphotericin B resistance in our isolate, it is important to note that reduced membrane lipid permeability and overexpression of *ERG* genes may have been contributors [1, 5, 19]. Finally, it is important to note that clade I isolates also have the highest rates of multidrug resistance (45%) and was the only clade to have extensive drug resistance (3%) to all three major classes of antifungals, as is the case with this clade I isolate [21].

A striking morphological feature of some *C. auris* isolates is their capacity to aggregate and form strong biofilms, an important risk factor for systemic infections [8, 11]. Our isolates differed significantly in their abilities to form biofilm and to aggregate, with the last isolate forming the strongest biofilm. Aggregation in clade I isolates was shown to be induced *in vitro* by exposure to antifungals likely due to cell wall changes [22]. Therefore, it is possible that aggregate formation typical of clade I isolates may contribute to the observed elevated antifungal MICs in this clade. Interestingly, the most highly aggregative isolate from our patient was the first one recovered following initiation of therapy. Although speculative, it is possible that aggregation could constitute a protective response induced by therapy to impede drug penetration into the cell aggregates. Although not within the scope of this study, it would be interesting to analyze the transcriptomes of these isolates for changes in expression of cell wall adhesins we recently showed to mediate aggregation and biofilm formation [11].

To assess whether development of resistance impacts cellular fitness, resistance to environmental stressors including thermotolerance was evaluated. Although the responses among the isolates varied, the terminal MDR isolate exhibited a notable slower growth rate, which has been proposed as a potential fitness cost for drug resistance development [13]. Another notable difference was the increased sensitivity of isolate #1 to the chitin-binding dye calcofluor white, which has been associated with aggregation, an interesting observation as this isolate was highly aggregative [23]. Although significance is not clear, these findings suggest potential differences in cell walls among the isolates, highlighting *C. auris* phenotypic plasticity.

The presence of both resistant and susceptible isolates in the same populations indicate that the resistance in *C. auris* is not intrinsic. With multi-drug resistance in *C. auris* on the rise and evolution of resistance closely linked to drug exposure, antifungals should be prescribed judiciously. Furthermore, given our findings demonstrating phenotypic variation among isolates in adhesion and cell-cell interaction, it would be interesting to explore whether evolved resistant strains have higher pathogenic potential. To our knowledge, this is the first study demonstrating *C. auris* evolution of MDR development in the same patient during therapy, underscoring the alarming rising threat of *C. auris*, further warranting accurate monitoring and surveillance of high-risk patients.

## ACKNOWLEDGEMENTS

The work in this publication was supported by the National Institute of Allergy and Infectious Diseases of the NIH under award number R01AI130170 (NIAID) to M.A.J-R. This work was also supported by the University of Maryland Baltimore, Institute for Clinical & Translational Research (ICTR). We would like to thank Indira French and Gwendolyn Paszkiewicz for assistance with antifungal susceptibility testing.

## AUTHOR CONTRIBUTIONS

M.A.J-R., T.W.W., N.P., J.K.J. conceived and designed this research, M.A.J-R. provided funding; T.W.W. performed experiments; M.A.J-R., T.W.W., N.P. analyzed data; M.A.J-R., T.W.W., N.P., J.K.J. wrote the paper; M.A.J-R. oversaw the study. All authors read and approved the manuscript.

## FIGURE LEGEND

**Figure 1. Timeline for isolation of *C. auris* clinical isolates, antifungal treatment and development of resistance. (A)** Patient was admitted for transplant evaluation with subsequent bilateral orthotopic lung transplant (BOLT). Timeline shows *C. auris* isolates from various timepoints and locations throughout hospitalization. Antifungals for treatment (orange), MICA: micafungin; IV LAMB: intravenous liposomal amphotericin B. Antifungals for transplant prophylaxis (grey), iLAMB: inhaled amphotericin B; ISA: isavuconazole. Isolate 4 resistant to caspofungin. Isolate 5 resistant to amphotericin B, caspofungin and micafungin. ETT, endotracheal tube; L-AMB, liposomal amphotericin **(B) Clinical antifungal susceptibilities of *C. auris* isolates**. MICs shown in bold exceed tentative CDC MIC (μg/ml) breakpoints for resistance: fluconazole (FZ) ≥32; 5-flucytocine (FC) >8; amphotericin B (AMB) ≥2; caspofungin (CAS) ≥2; rezafungin (RZF) S ≤0.5; voriconazole (VOR) S ≤0.12; itraconazole (IZ) ≤0.125; micafungin (MF) ≥4; (https://www.cdc.gov/candida-auris/hcp/laboratories/antifungal-susceptibility-testing.html). There is no CDC advisory for FZ, IZ and VOR breakpoints; MICs are considered resistant based on breakpoints for other *Candida* species

**Figure 2. Comparative phenotypic evaluation of clinical isolates. (A)** Evaluation of cell growth rates among isolates over 36h based on values of OD_600_. Line plots show mean and standard error of 10 technical replicates. **(B)** Measurement of metabolic activity of 24h biofilms based on values of OD_490_. Bar-plots show mean and standard error of mean of *n* = 3 biological replicates, each as an average of 4 technical replicates. **(C)** Measurement of rate of cell aggregate sedimentation over 30 mins. Bar-plots show mean (n = 3) of biological replicates, each an average of 3 technical replicates. **(D)** Representative light microscopy (10x) images of cell aggregates formed by isolates. Statistical analysis was performed by one-way ANOVA and post-hoc Tukey test with *P*-values representing significant differences. *0.01 < P ≤ 0.05, **0.001 < P ≤ 0.01, ***0.0001 < P ≤ 0.001. Non-significant differences are not shown.

**Supplemental Figure 1. Response of clinical isolates to environmental stresses. (A)** Heat maps depicting percent change in growth relative to control in the presence of SDS, H_2_O_2_, and NaCl as an average of 2 biological replicates. **(B)** Isolates grown in presence of calcofluor white (CW) (10mg/L) and at various growth temperatures and cell densities (1×10^6^,1×10^5^,1×10^4^ cells/ml).

**Supplemental Table 1.** Primers used in this study.

