## Supplemental Table 1 for "*In vivo* evolution of *Candida auris* multi-drug resistance in a patient receiving antifungal treatment"

**Supp Table 1. Primers used in this study**

| Gene | Primer Sequence 5'- 3' |
|---|---|
| *ITS1* | TCCGTAGGTGAACCTGCGG |
| *ITS1* CAU-R | TTTGTGAATGCAACGCCATCG |
| *RHA1-F* | TTGCGGTTGAAATGGGTGCT |
| *RHA1-R* | TGGCATGTTTCCGGCTTAGA |
| *ERG11-F* | TCGCGTAAATACAATGCCC |
| *ERG11-R* | TGGTTTGGTGAAGAATTCGG |
| *ERG6-F* | CACTTCAACTGTTCCAAC |
| *ERG6-R* | GGTTATCGTGAATGGCAT |
| *FKS1* HSP1-F | GCCATCTCGAAGTCTGCTCA |
| *FKS1* HSP1-R | TGACAATGGCATTCCACACCT |
| *FKS1* HSP2-F | GCGAGAACCTTGGCTCAAA |
| *FKS1* HSP2-R | ATGGCAAGAAGTCAGCCATGA |
