## Supplemental Figure 1 for "*In vivo* evolution of *Candida auris* multi-drug resistance in a patient receiving antifungal treatment"

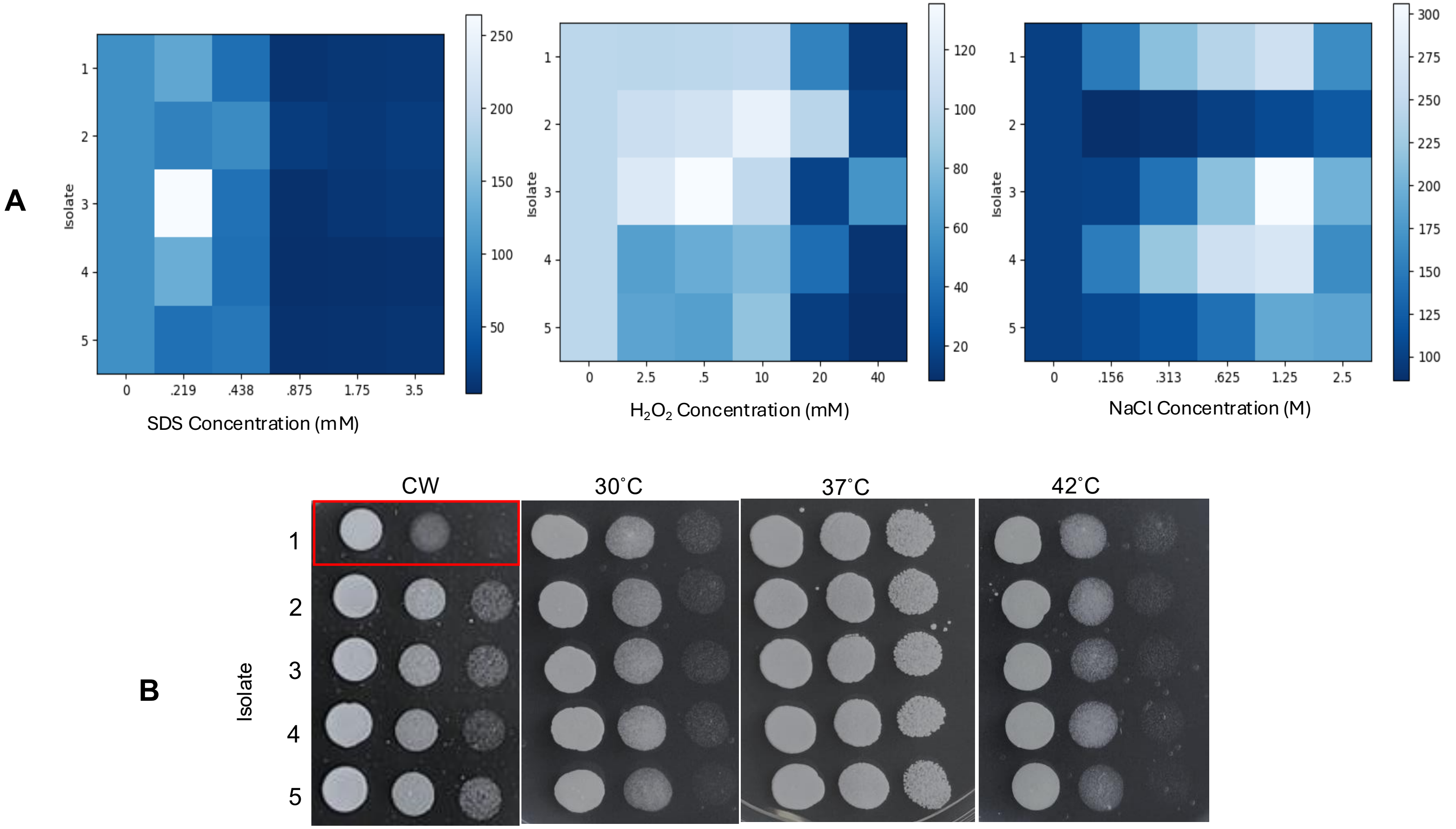


**Supplemental Figure 1. Response of clinical isolates to various stresses. (A)** Heat maps depicting percent change in growth relative to control in the presence of SDS, $H_2O_2$, and NaCl as an average of 2 biological replicates. **(B)** Isolates grown in presence of calcofluor white (CW) (10mg/L) and at various growth temperatures and cell densities ($1{\times}10^6$,$1{\times}10^5$,$1{\times}10^4$ cells/ml).
